# Evaluation of Locomotor Activity in Group-Housed Hamsters: Effects of Cage Size and Running Wheel Availability

**DOI:** 10.64898/2026.08.24.746863

**Authors:** Junya Sato, Ayato Mase, Mizuki Ito, Keishirou Yoshida

## Abstract

While hamsters are commonly housed in groups within pet shops in Japan, small cages are often thought to restrict their physical activity, leading to arguments that larger cages should be provided. Although previous research has investigated how cage size and running wheel availability influence activity levels in individually housed hamsters, no studies to date have examined these specific effects in a social housing context. Therefore, this study investigated how cage size and the presence of a running wheel affect the activity levels of individual hamsters using the group-housing conditions with five hamsters. Video recordings were captured for 24 hours across four distinct cage environments using a camera installed directly above each cage. From these recordings, the distances traveled on both the cage floor and the running wheel were calculated for each hamster and statistically analyzed. The results revealed that overall activity levels were significantly higher in cages equipped with a running wheel than in those without. Although cage size did not yield a statistically significant difference, a marginal trend toward higher activity in larger cages was observed.

## 1 Introduction

Hamsters are among the most popular pets and are housed in various facilities, including pet shops. Due to space and cost limitations in commercial facilities, hamsters are commonly maintained in group-housed environments. However, because they are naturally solitary and highly territorial, this housing practice raises critical welfare concerns; restricting inherently aggressive and territorial rodents to small, shared cages often leads to decreased physical activity and elevated stress levels.

Numerous studies have investigated the effects of cage size or running wheel availability on a hamster under solitary housing conditions. For instance, Fischer *et al*. examined how activity levels varied across four different cage sizes [1]. Although no significant differences in activity were found, the hamsters gnawed at the wire for longer periods in smaller cages. Similarly, Kuhnen suggested that housing in confined spaces potentially induces stress [2]. The effect of the presence of a running wheel has also been extensively documented. Gebhardt-Henrich *et al*. compared the effects of functional versus non-functional running wheels, finding that individuals provided with functional wheels exhibited significantly fewer stereotypic behaviors, such as wire-gnawing and cage-climbing [3]. Gattermann *et al*. conducted a long-term comparison between groups with and without access to running wheels; their results showed that the group with access to running wheels experienced greater increases in body weight and food intake, highlighting the importance of a running wheel for animal welfare [4]. Furthermore, Reebs *et al*. used preference tests to investigate favored wheel sizes, shapes, and materials to determine ideal specifications from a welfare perspective [5]. Regarding wheel design, Mrosovsky *et al*. compared activity levels in animals using wheels with plastic mesh surfaces and those with standard metal rungs [6]. Finally, Scherbarth *et al*. and Kingston *et al*. investigated how voluntary wheel running affects seasonal adaptation for winter and how it enhances resilience to the stress induced by social defeat, respectively [7, 8].

As described by these studies, the effects of cage size and availability of a running wheel on hamster activity levels are typically analyzed within solitary housing environments, probably due to the difficulty of measuring individual activity levels of group-housed hamsters. For instance, when multiple hamsters share a single running wheel, a method capable of identifying which individual is using the wheel and for how long is required. To the best of our knowledge, no previous study has established such a method. Consequently, it remains unclear to what extent cage size and the presence of a running wheel influence the activity levels of individual hamsters under group housing conditions. To address this gap, five golden hamsters (*Mesocricetus auratus*) were filmed over a 24-hour period in four distinct housing environments (large or small cages and with or without a running wheel). Using the recorded videos, we statistically analyzed the activity levels of each hamster on both the cage floor and running wheel.

## 2 Experimental Setup

Two cage sizes were used in this study, a large cage measuring 580 × 392 × 400 mm and a small cage measuring 368 × 222 × 262 mm (Fig. 1). The same five hamsters were observed in each cage size, with and without a running wheel, resulting in the four housing conditions/treatments. The running wheel had a diameter and circumference of 170 and 533 mm, respectively. To visually measure the number of rotations, a black rectangular marker was attached to the wheel, as indicated by the red circle in the upper-left of Fig. 1. Since visually identifying and continuously tracking hamsters with similar appearances is challenging, five golden hamsters, *Mesocricetus auratus*, with distinct coat patterns were used. All hamsters were females aged three months or older and standardized at approximately four months of age, with a head-body length of 10 cm. The sides of the cages used were made of glass, and the trays were composed of polystyrene. The experiment was conducted in a constant-temperature room maintained at 24°C, where the indoor light/dark cycle depended on natural ambient light. To capture the hamsters’ activity during both day and night, an infrared camera was installed overlooking the cage. The recorded video resolution was 800 × 448 pixels. During our visual inspections, no injuries resulting from hamsters getting caught in the wheel were observed, indicating that the setup did not pose a physical hazard.

**Fig. 1:**
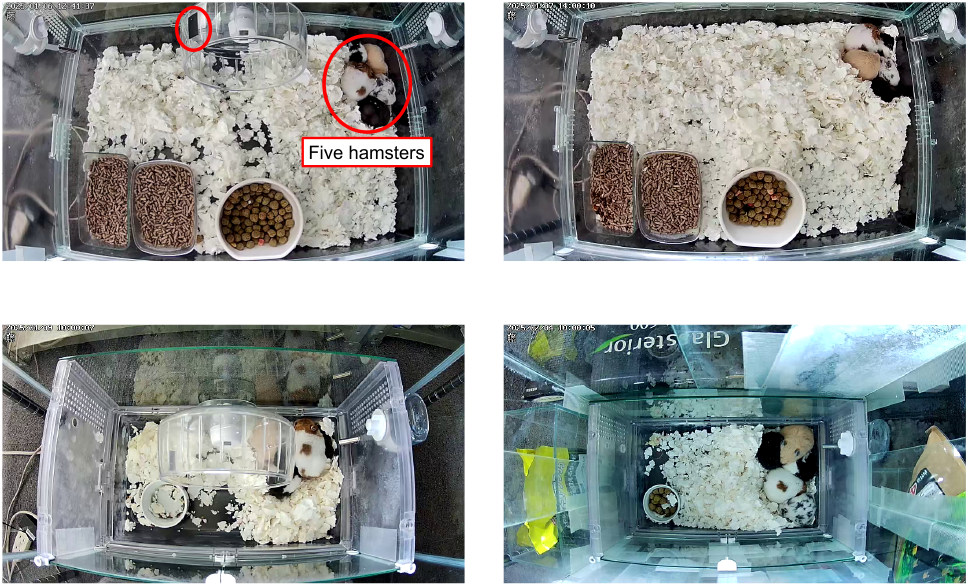
Top row: 580 × 392 × 400 mm large cages; bottom row: 368 × 222 × 262 mm small cages. Left column: with running wheel; right column: without running wheel. The black mark circled in red in the top-left is attached to visually measure the number of rotations.

## 3 Measurement of Traveled Distance

The distance traveled by each hamster using the running wheel was calculated by visually counting the number of rotations based on the black markers attached to the wheel and multiplying that count by the circumference (533 mm). For the distance traveled on the cage floor, measurements were taken after applying image processing. First, using background subtraction, all frames in which moving objects appeared were saved. Fig. 2 shows an example of this process. This preprocessing allowed us to exclude a large number of frames where no movement was detected.

**Fig. 2:**
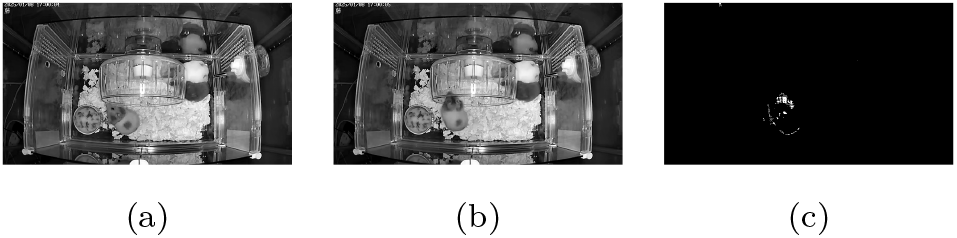
Extraction of a frame containing moving objects using background subtraction: (a) previous frame; (b) current frame; (c) background subtraction result.

In all extracted frames, the head of each hamster was annotated with a point using LabelMe [9]. Fig. 3 illustrates an example of an annotation for a single individual. The total floor distance was calculated by integrating the Euclidean distances between consecutive annotated points. Since the initial measurements were in pixels, the change to meters was achieved by determining the ratio between the actual cage dimensions and their corresponding size within the images.

**Fig. 3:**
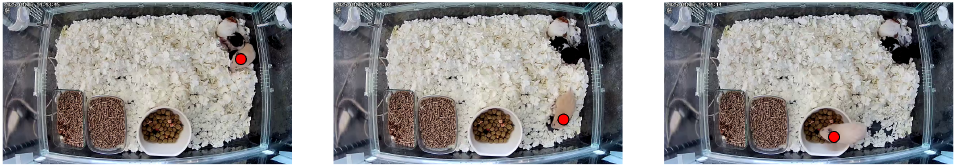
Examples of an annotation (red circle).

## 4 Results and Discussion

The measured travel distances per hamster are presented in Fig. 4. As shown in (a), the distance traveled on the running wheel exceeded that on the cage floor in both cage sizes. The fact that hamsters devote a significant portion of their active time to wheel-running is consistent with the findings of Fischer *et al*. [1]; our results demonstrate that this behavioral pattern persists even in group-housed environments.

**Fig. 4:**
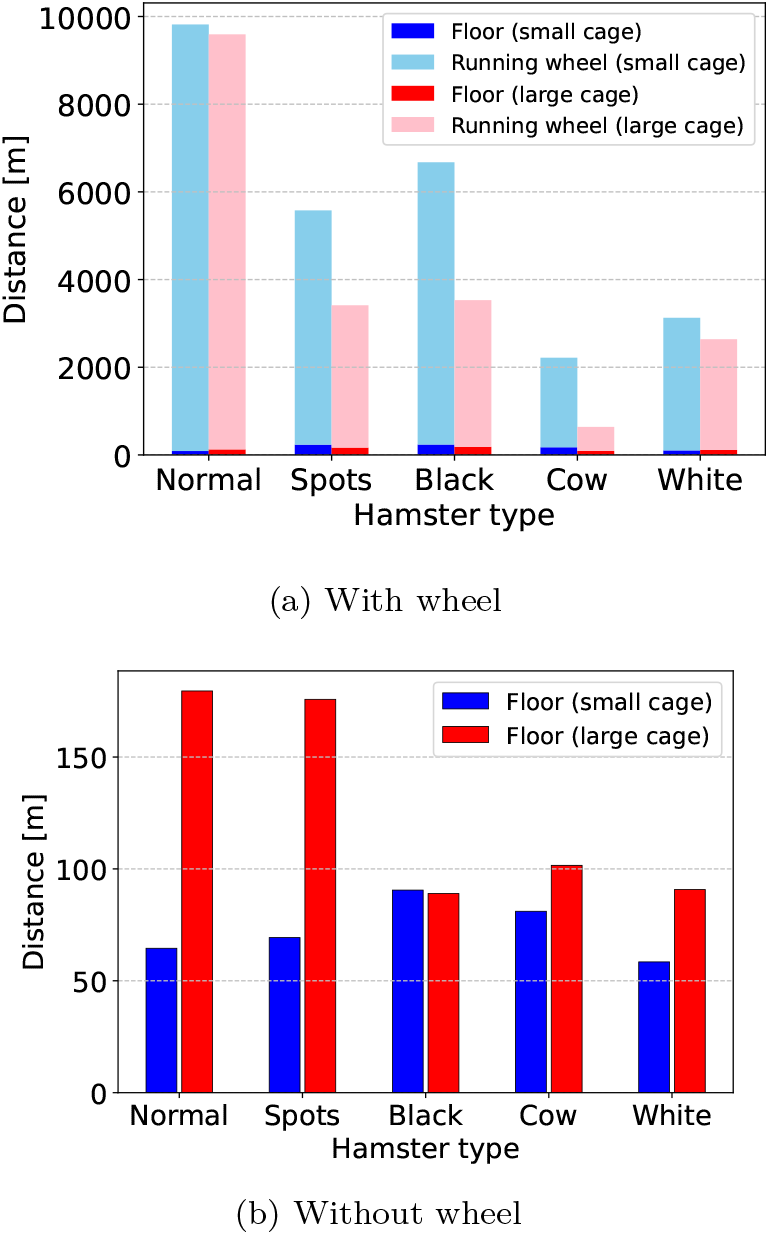
Locomotor distance of hamsters.

The total travel distance per individual was higher in the smaller cages than that in the larger ones. Contrary to our initial hypothesis, this may be attributed to an increased reliance on the running wheel as a means of stress overcompensation due to the confined space. Furthermore, the result for cages without running wheel (Fig. 4(b)) showed greater travel distances in the larger cages, suggesting that hamsters may have spent more time engaged in exploratory behavior within the increased floor area.

Comparing the relationship between the presence of running wheel and the traveled distance in Fig. 4(a) and (b), it is evident that activity levels are significantly higher when a wheel is available. Considering that the average daily travel distance for a hamster is approximately 9 km [1, 4, 10], the provision of a running wheel is arguably essential. In our group-housed environment, however, only one individual (”normal”) exceeded this average distance. Because hamsters prefer a solitary lifestyle and are prone to aggression when group-housed, the competition for the running wheel may have been driven by social dominance hierarchies. Although we did not assess agonistic behaviors in the present study, our next research will incorporate behavioral observations of aggression to analyze its relationship with locomotor distance.

Fig. 5 shows a large dispersion in the total travel distance among individuals. This suggests that under social housing conditions, inequality in access to the running wheel may occur. Observations of the recorded video revealed instances where multiple hamsters attempted to simultaneously use the wheel as shown in Fig. 6. Therefore, installing multiple running wheels might alleviate this inequality. While selecting five golden hamsters with distinct coat patterns was necessary to facilitate visual tracking and individual identification during video analysis, this choice introduces a potential source of variation. Phenotypic differences could inherently influence the group’s social structure, hierarchical dominance, and individual access to shared resources. In this study, we did not formally assess whether specific individuals exhibited social dominance, actively displaced conspecifics, or monopolized the single running wheel. Given the small number of animals used, the lack of a direct evaluation of these phenotypic and social hierarchy effects must be acknowledged as a limitation. Future studies should incorporate formal dominance scoring to clarify how individual social status interacts with resource utilization in group-housed conditions. The results of the two-way repeated measures ANOVA are listed in Table 1. There was a significant main effect of the presence of a running wheel (*F* (1, 4) = 10.98, *p* = .030) and a significant interaction between cage size and the presence of running wheel (*F* (1, 4) = 8.89, *p* = .041). In contrast, while the main effect of cage size was not statistically significant, a marginal trend toward significance was observed (*F* (1, 4) = 7.12, *p* = .056). These results suggest that the impact of installing a running wheel on the distance traveled varies depending on the size of the cage.

**Table 1:** Results of two-way repeated measures ANOVA for hamster locomotor distance. Note that *df* = degrees of freedom (numerator, denominator) and * significant at the .05 level.

| Source | $df$ | $F$ | $p$ |
| --- | --- | --- | --- |
| Cage size (Size) | 1, 4 | 7.12 | .056 |
| Running wheel (Wheel) | 1, 4 | 10.98 | .030* |
| Size × Wheel | 1, 4 | 8.89 | .041* |

**Fig. 5:**
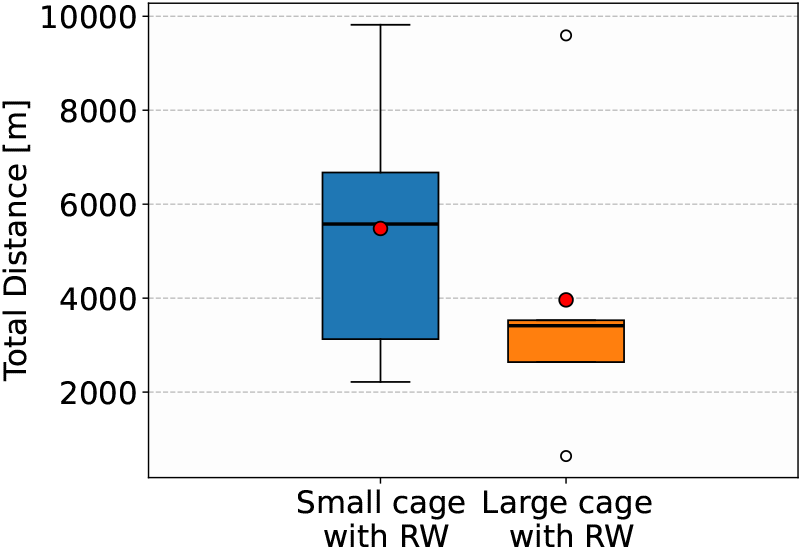
Box-and-whisker plots based on the distance traveled by hamsters in large and small cages equipped with a running wheel.

**Fig. 6:**
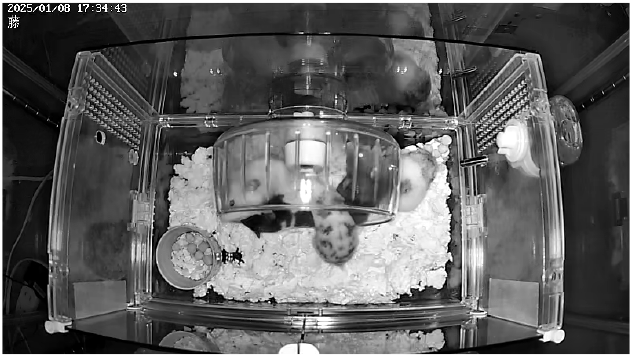
Four hamsters trying to use the running wheel.

Since the interaction effect was significant, the simple main effect of wheel availability for each cage size (Table 2) was examined. In the small cage, the presence of a running wheel significantly increased the locomotion distance (*t*(4) = 4.01, *p* = .016). In contrast, while an increasing trend was observed in the large cage, the difference did not reach statistical significance (*t*(4) = 2.58, *p* = .061). These results suggest that in a small cage, a running wheel is highly effective in promoting locomotor activity. The observed difference between wheel conditions may have been less pronounced in the large cage, as the increased floor area allowed for more exploratory behavior. Consequently, providing a running wheel appears particularly important when floor space is restricted, serving as a valuable tool to facilitate exercise opportunities.

**Table 2:**
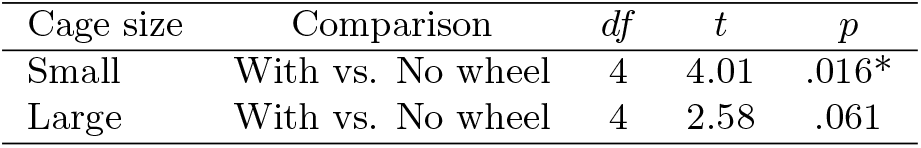
Simple main effect analysis of running wheel availability across different cage sizes. Note that *df* = 4 based on the number of subjects (*n ™* 1) and * significant at the .05 level.

## 5 Conclusion

This study investigated the effects of cage size and the availability of a running wheel on the activity levels of individual hamsters under group-housed conditions. The results identified that the provision of a running wheel significantly increased the activity levels of hamsters, with this effect being particularly pronounced in the small cage condition. Furthermore, a significant interaction was observed between cage size and wheel availability, suggesting that running wheels may be particularly important for facilitating increased locomotor activity in small cages. However, it remains unclear whether this increased activity solely reflects a motivation for exercise. Alternatively, the high wheel-running activity in confined spaces could be driven by other behavioral or physiological mechanisms, such as a lack of exploratory space, frustration, or stress-coping behaviors. For instance, wheel running has been shown to attenuate stress responses in rodents [11]. Further investigation is required to determine the underlying motivational and emotional processes driving this behavior in smaller housing environments. Additionally, we observed instances where multiple hamsters competed for a single wheel, alongside individual differences in locomotion levels. Therefore, future research should examine whether increasing the number of running wheels can mitigate these individual disparities and improve the overall activity levels of the group.

## Acknowledgements

We would like to express our sincere gratitude to Dr. Osamu Tanaka of Kuu Animal Hospital for his invaluable advice regarding the ecology of golden hamsters.

## Statements and Declarations

### Funding

This study was supported by Japan Bird & Small Animal Association.

### Conflict of interest

The authors declare no conflict of interest.

### Ethics approval

This study was conceived and conducted by GEX Co., Ltd. All experiments were performed in accordance with the Act on Welfare and Management of Animals established by the Ministry of the Environment of Japan. We ensured maximum consideration for animal welfare by following the Guidelines for Proper Conduct of Animal Experiments by the Science Council of Japan, the notices of the Ministry of Education, Culture, Sports, Science and Technology (MEXT), and the guidelines for animal experiments provided by the Japanese Association for Laboratory Animal Science. The principal investigator evaluated the ethical validity of the experimental design in light of these official guidelines beforehand, and the 3Rs principles (Replacement, Reduction, and Refinement) were strictly maintained throughout the entire process.

### Data availability

The datasets are available from the corresponding author on reasonable request.

### Author contribution

All authors contributed to the study conception and design. Material preparation and video collection were performed by Keishirou Yoshida. The analysis was performed by Mizuki Ito, Ayato Mase and Junya Sato. The first draft of the manuscript was written by Junya Sato. The review and editing was performed by Keishirou Yoshida.

## Notes

### Competing Interest Statement

The authors have declared no competing interest.

